# FIP200 organizes the autophagy machinery at p62-ubiquitin condensates beyond activation of the ULK1 kinase

**DOI:** 10.1101/2020.07.07.191189

**Authors:** Eleonora Turco, Irmgard Fischer, Sascha Martens

## Abstract

Macroautophagy is a conserved degradation pathway, which mediates cellular homeostasis by the delivery of harmful substances into lysosomes. This is achieved by the sequestration of these substances referred to as cargo within double membrane vesicles, the autophagosomes, which form *de novo*. Among the many cargoes that are targeted by autophagy are condensates containing p62 and ubiquitinated proteins. p62 recruits the FIP200 protein to initiate autophagosome formation at the condensates. How FIP200 in turn organizes the autophagy machinery is unclear. Here we show that FIP200 is dispensable for the recruitment of the upstream autophagy machinery to the condensates, but it is necessary for phosphatidylinositol 3-phosphate formation and WIPI2 recruitment. We further find that FIP200 is required for the activation of the ULK1 kinase. Surprisingly, ULK1 kinase activity is not strictly required for autophagosome formation at p62 condensates. Super-resolution microscopy of p62 condensates revealed that FIP200 surrounds the condensates where it spatially organizes ATG13 and ATG9A for productive autophagosome formation. Our data provide a mechanistic insight into how FIP200 orchestrates autophagosome initiation at the cargo.

## Introduction

Macroautophagy, hereafter autophagy, is a degradative pathway that contributes to the maintenance of cell homeostasis and survival, through the delivery of harmful substances into lysosomes for degradation. Among the many substances, referred to as cargo, that are degraded by autophagy are damaged organelles, pathogens and protein aggregates. Misfunctions of autophagy are associated with a plethora of diseases including neurodegeneration and cancer (Levine & Kroemer, 2019).

Upon induction of autophagy, cells respond with the assembly of a membrane structure termed isolation membrane (or phagophore). The isolation membrane expands and captures cytoplasmic cargo as it grows. Subsequently, the isolation membrane closes to give rise to a double membrane autophagosome, which fuses with a lysosome wherein the inner membrane and the cargo are degraded (Morishita & Mizushima, 2019).

Autophagosome formation can be triggered by inhibition of mTOR upon starvation (Alers, Loffler et al., 2012, Noda & Ohsumi, 1998) and also by the presence of cargo material in the cytoplasm, which signals its autophagic degradation via cargo receptors (Ravenhill, Boyle et al., 2019, Turco, Fracchiolla et al., 2020, Turco, Witt et al., 2019, Vargas, Wang et al., 2019, Zaffagnini & Martens, 2016). In both scenarios autophagosome formation requires the orchestrated action of the core autophagy machinery (Xie & Klionsky, 2007).

This machinery is composed of the ULK1 complex, containing the ULK1 kinase and the FIP200/RB1CC1, ATG13 and ATG101 subunits (Ganley, Lam du et al., 2009, Hara, Takamura et al., 2008, Hosokawa, Hara et al., 2009, Jung, Jun et al., 2009, Mercer, Kaliappan et al., 2009, Yan, Kuroyanagi et al., 1998), Golgi derived vesicles containing the ATG9A protein (Young, 2006), the class III phosphatidylinositol-3 kinase complex 1 (PI3KC3-C1) which converts phosphatidylinositol (PI) to phosphatidylinositol 3-phosphate (PI3P) (Itakura & Mizushima, 2009, Matsunaga, Saitoh et al., 2009, Obara, Sekito et al., 2006), the PI3P binding WIPI proteins (Dooley, Razi et al., 2014, Proikas-Cezanne, Takacs et al., 2015), the ATG2 lipid transfer proteins (Valverde, Yu et al., 2019) as well as the ATG12 and LC3 conjugation systems (Ichimura, Kirisako et al., 2000, Martens & Fracchiolla, 2020, Mizushima, Noda et al., 1998). In starvation induced autophagy, these components act in a hierarchical manner (Itakura & Mizushima, 2010, Koyama-Honda, Itakura et al., 2013). Loss of mTOR signaling results in localization and activation of the ULK1 complex at ER proximal sites, followed by ATG9 vesicles and PI3KC3-C1 recruitment, which in turn attract the WIPI and ATG2 proteins. This is followed by recruitment of the ATG12–ATG5-ATG16L1 complex (hereafter E3), which acts analogous to E3 complexes in ubiquitination reactions in the subsequent conjugation of LC3 proteins to isolation membrane resident phosphatidylethanolamine (PE), a reaction known as lipidation (Hurley & Young, 2017, Martens & Fracchiolla, 2020, Melia, Lystad et al., 2020).

During cargo-induced selective autophagy, which can occur in the presence of mTOR signaling, cargo receptors trigger the formation of an isolation membrane around the cargo by recruiting the ULK1 complex subunit FIP200 (Ravenhill et al., 2019, Smith, Harley et al., 2018, Turco et al., 2019, Vargas et al., 2019). The precise mechanism by which cargo receptor mediated FIP200 recruitment initiates autophagosome formation at the cargo remains to be elucidated.

FIP200 is essential for both starvation-induced and selective autophagy (Hara et al., 2008, Kishi-Itakura, Koyama-Honda et al., 2014). In starvation-induced autophagy, FIP200 was shown to localize at the ER, potentially bridging the autophagy machinery with the ER scaffold and it appeared as a cup-shaped structure by super-resolution microscopy (Kenny, Chen et al., 2019). Recently, electron microscopy analysis of purified ULK1-ATG13-FIP200 complex, revealed that the N-terminal domain of FIP200 forms a C-shaped scaffold which organizes the other subunits of the complex (Shi, Yokom et al., 2020).

Here, we studied the selective autophagy process known as aggrephagy, mediated by the cargo receptor p62/SQSTM1. p62 phase separates ubiquitinated proteins into larger condensates to promote their degradation (Bjorkoy, Lamark et al., 2005, Danieli & Martens, 2018, Sun, Wu et al., 2018, Zaffagnini, Savova et al., 2018). We previously discovered that p62 directly interacts with a C-terminal Claw domain of FIP200 and this interaction mediates the autophagic degradation of p62 condensates (Turco et al., 2019). How the recruitment of FIP200 to the condensates initiates their degradation is unclear.

Here, we show that the role of FIP200 in the selective autophagy of p62-ubiquitin condensates goes beyond the recruitment of the upstream autophagy machinery. In fact, FIP200 is not strictly required for the recruitment of the ULK1 complex and the PI3KC3-C1 to the condensates. However, it is necessary for the activation of both kinase complexes, for WIPI2 and E3 recruitment and LC3 lipidation. Surprisingly, we find FIP200 to be essential for the progression of autophagosome formation, while ULK1 activity is dispensable. Finally, we observe FIP200 to form cup-shaped structures around p62 condensates and find that it is required for the correct positioning of ATG13 and ATG9A vesicles. We propose that it acts as a scaffold for the correct spatial organization of the autophagy initiation machinery, leading to PI3P production, likely on ATG9A vesicles, and subsequently autophagosome formation around the cargo.

## Results and discussion

### FIP200 is essential for the clearance of p62 condensates by selective autophagy

To study the role of FIP200 in selective autophagy we employed HAP1 cells cultivated in nutrient rich medium and we compared wild type (WT) to FIP200 KO cells. In absence of external autophagy stimuli such as starvation, cells perform autophagy at basal levels to maintain homeostasis, which in the case of aggrephagy can be monitored by following the levels of p62 and lipidated LC3B (LC3B-II). In FIP200 KO cells LC3B lipidation is abolished and a strong accumulation of p62 in comparison to WT cells is observed. Inhibition of lysosome acidification by Bafilomycin A1 treatment caused no increases in LC3B-II levels, suggesting that FIP200 deletion blocks autophagy (Figure 1A). This observation is in accordance with previous reports showing that FIP200 is essential for both starvation induced and selective autophagy (Hara et al., 2008, Itakura & Mizushima, 2010, Ravenhill et al., 2019, Turco et al., 2019, Vargas et al., 2019). Immunofluorescence staining of p62 in FIP200 KO cells showed a higher number of p62 condensates with a 4,5-fold increase in their area when compared to WT cells (Figure 1B), consistent with a block in their autophagic turnover. FIP200 is recruited to p62 condensates through direct interaction of its Claw domain with p62 and deletion of the entire Claw domain impairs autophagy (Turco et al., 2019). Since FIP200 Claw domain may have other autophagy related functions, here we took a more specific approach to assess the role of p62-FIP200 interaction in the selective autophagy of p62-ubiquitin condensates. Using CRISPR/Cas9 we generated a cell line in which the arginine 1573 of endogenous FIP200 was mutated into aspartate. This R1573D point mutation was shown to abolish the interaction of FIP200 with p62 *in vitro* (Turco et al., 2019). The point mutation did not affect FIP200 expression levels and resulted in an elevated level of the p62 protein and reduced LC3B lipidation (Figure 1A), suggesting that Claw domain-mediated recruitment of FIP200 to cargo material is responsible for a considerable fraction of the basal autophagic activity in the cells. In contrast, upon induction of autophagy by starvation the difference between the FIP200 WT and R1573D cells in the levels of lysosomal delivered LC3B-II was much smaller, pointing to a specific function of the Claw domain in selective autophagy (Figure EV1).

**Figure 1.**
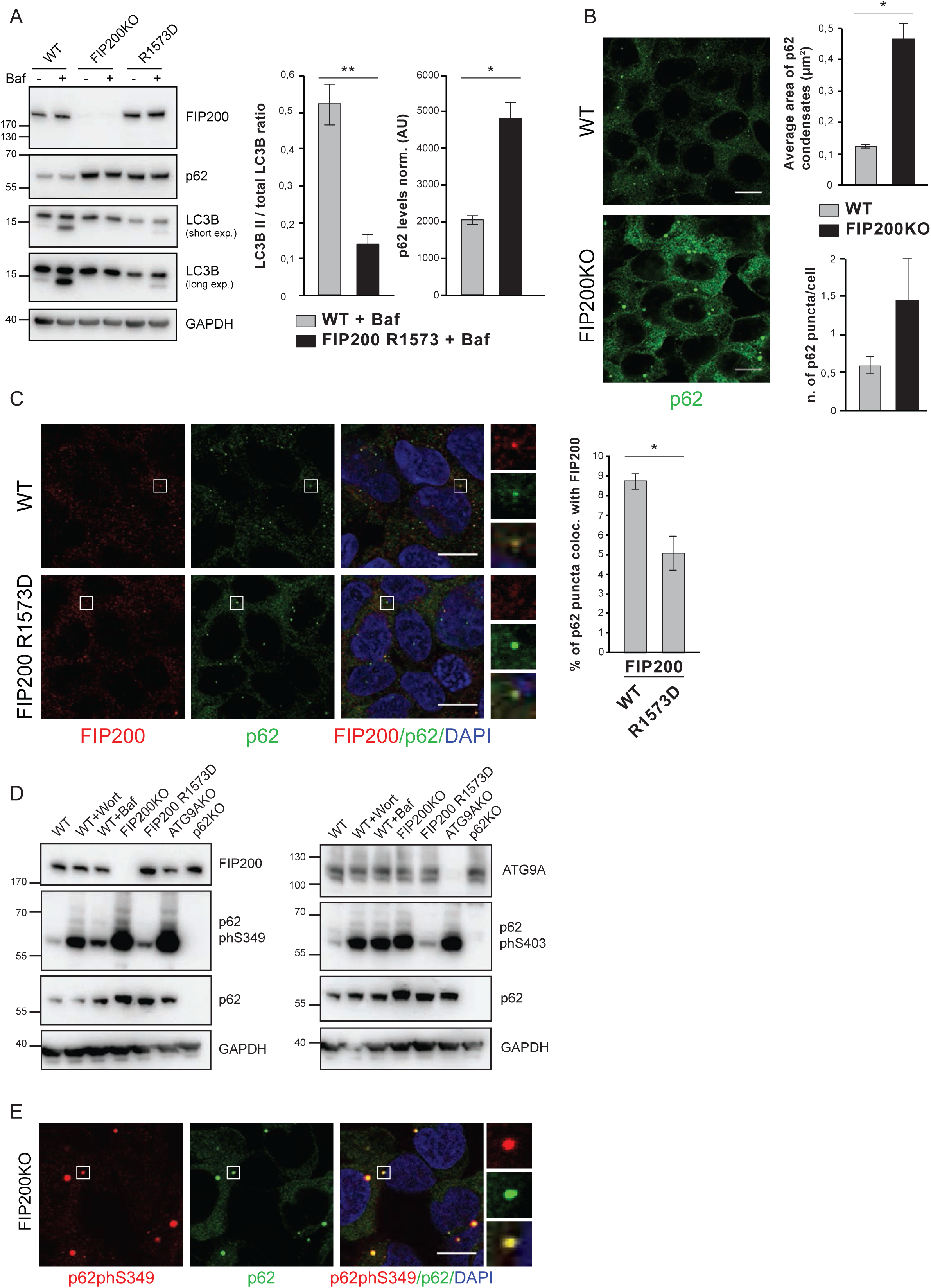
FIP200 is essential for clearance of p62 condensates by selective autophagy. (A) HAP 1 cells (WT, FIP200 KO or FIP200 R1573D) were left untreated or treated with 400 nM Bafilomycin A1 for 2 h. Cell lysates were analyzed by western blotting with anti-FIP200, anti-p62 and anti-LC3B antibodies. The LC3B-II/LC3B-I ratio and the levels of p62 in Bafylomycin A1 treated cells are plotted on the right. Mean and S.E.M. are shown for n=3. Significances are marked with * when p value ≤ 0,05, ** when p value ≤ 0,01, *** when p value ≤ 0,001. (B) Analysis of p62 condensates in HAP1 WT and FIP200 KO cells. Endogenous p62 was detected by immunofluorescence. Scale bar = 10 µm. The number of p62 puncta and their area (µm^2^) is plotted on the right. Mean values and S.E.M. are shown for n=3. Significances are marked with * when p value ≤ 0,05, ** when p value ≤ 0,01, *** when p value ≤ 0,001. (C) Colocalization analysis of p62 and FIP200 in HAP 1 WT and FIP200 R1573D cells. p62 and FIP200 were detected by immunofluorescence. White sboxes represent the magnified areas shown on the right. Scale bar = 10 µm. The percent of p62 puncta colocalizing with FIP200 is plotted on the right. Average percentages of colocalization and S.E.M are shown for n=3. Significantly different samples are marked with * when p value ≤ 0,05, ** when p value ≤ 0,01, *** when p value ≤ 0,001. (D) Analysis of phosphorylated p62 levels in HAP 1 cell lines. WT or the mutant cell lines were left untreated or treated with Wortmannin (1 µM for 2 h) or Bafilomycin A1 (400 nM for 2 h). Cell lysates were analyzed by western blotting. FIP200, ATG9A and p62 antibodies were used as a control for the knock out cell lines. Levels of phosphorylated p62 were analyzed with anti-p62 phS349 (left) and anti-p62 phS403 (right) antibodies. (E) p62 and p62phS349 were detected by immunofluorescence in HAP 1 FIP200 KO cells. Scale bar = 10 µm. White boxes indicate the magnified areas shown on the right.

FIP200 R1573D cells also showed a significantly reduced recruitment of FIP200 to p62 condensates (Figure 1C). The residual colocalization between FIP200 and p62 in FIP200 R1573D mutant cells, as well as the residual LC3B lipidation may be due to the presence of other cargo receptors or alternative recruitment mechanisms. For instance TAX1BP1, a cargo receptor of the CALCOO gene family, was proposed to have a role in aggrephagy (Sarraf, Shah et al., 2019) and was shown to interact with FIP200 outside the Claw domain (Ravenhill et al., 2019). Additionally, other autophagy factors such as ATG9A may mediate ULK1 complex and thus FIP200 recruitment.

These data suggest that the Claw domain specifically mediates the recruitment of FIP200 to p62-ubiquitin condensates in cargo-induced selective autophagy. This is further corroborated by the fact that the p62 phosphorylated at S349, which binds stronger to the Claw domain (Turco et al., 2019), dramatically accumulates in FIP200 KO cells and to some extent also in FIP200 R1573D cells (Figure 1D, E). This is consistent with previous reports, and suggests that this form of p62 is preferentially targeted by autophagy (Ichimura, Waguri et al., 2013) (Relic, Charlier et al., 2018). We observed a similar pattern for P-S403 p62 (Figure 1D), which was shown to display increased ubiquitin binding and p62-ubiquitin condensate formation (Matsumoto, Wada et al., 2011, Zaffagnini et al., 2018).

### Recruitment of autophagy machinery to p62 condensates

Next, we asked which steps of autophagosome formation at p62-ubiquitin condensates are perturbed in FIP200 KO cells by following the recruitment of the autophagy machinery to them. Due to the transient nature of some interactions and the high turnover of p62-positive autophagic structures in WT cells, we used the VPS34 inhibitor Wortmannin to accumulate early autophagy structures. We also used ATG9A KO cells to inhibit autophagy at a very early stage, just downstream of the ULK1 complex assembly (Karanasios, Walker et al., 2016, Orsi, Razi et al., 2012). As expected, analysis of ATG9A KO cells by immunoblot revealed that ATG9A is necessary for the clearance of protein aggregates as they showed a strong accumulation of P-S349 and P-S403 p62, similar to FIP200 KO cells (Figure 1D, EV2A). Lipidation of LC3B is almost completely abolished (Figure EV2A). Immunofluorescence staining of p62 in ATG9A KO cells showed accumulation of p62 condensates with a 4-fold increase in size when compared to WT cells (Figure EV2B).

We then proceeded to analyze the recruitment of the early autophagy machinery to p62 condensates by immunofluorescence. ATG13 partially colocalized with p62 in WT cells and the colocalization became more evident in Wortmannin treated cells (Figure 2A). Because both Wortmannin treatment and FIP200 deletion cause the accumulation of early autophagy structures, upstream of LC3B lipidation (Itakura & Mizushima, 2010), we compared ATG13 recruitment in FIP200 KO and Wortmannin treated cells. We observed that ATG13 is still recruited to p62 condensates in absence of FIP200, albeit less efficiently than in Wortmannin treated cells (*p value*: 0,059) and in ATG9 KO cells (Figure 2A). We observed that FIP200 aids to the recruitment of ULK1 complex to p62 condensates because when we knocked down FIP200 in ATG9 KO cells by siRNA (Figure EV2D) a strong reduction of ATG13 recruitment was observed. In addition, also the signal intensity of ATG13 puncta at the p62 condensates was reduced upon FIP200 depletion in ATG9A KO cells. (Figure 2B). We then observed a similar result for the recruitment of ULK1 to p62 condensates in absence of FIP200 (Figure EV2C). It was previously shown that deletion of FIP200 cause instability of ULK1 complex proteins (Hara et al., 2008). Indeed, we observed a reduction of ULK1 and ATG13 protein levels in FIP200 KO cells (or upon FIP200 knock down) (Figure EV2D), which could partly explain the decrease of ULK1 and ATG13 signal at p62 condensates. We conclude that FIP200 is a major contributor of ATG13 and ULK1 recruitment to p62-ubiquitin condensates. By contrast, we observed that ATG9A recruitment to p62 condensates is independent of FIP200 (Figure 3A). Similarly, ATG14 localization to p62 puncta was not affected by FIP200 or ATG9A deletion (Figure 3B).

**Figure 2.**
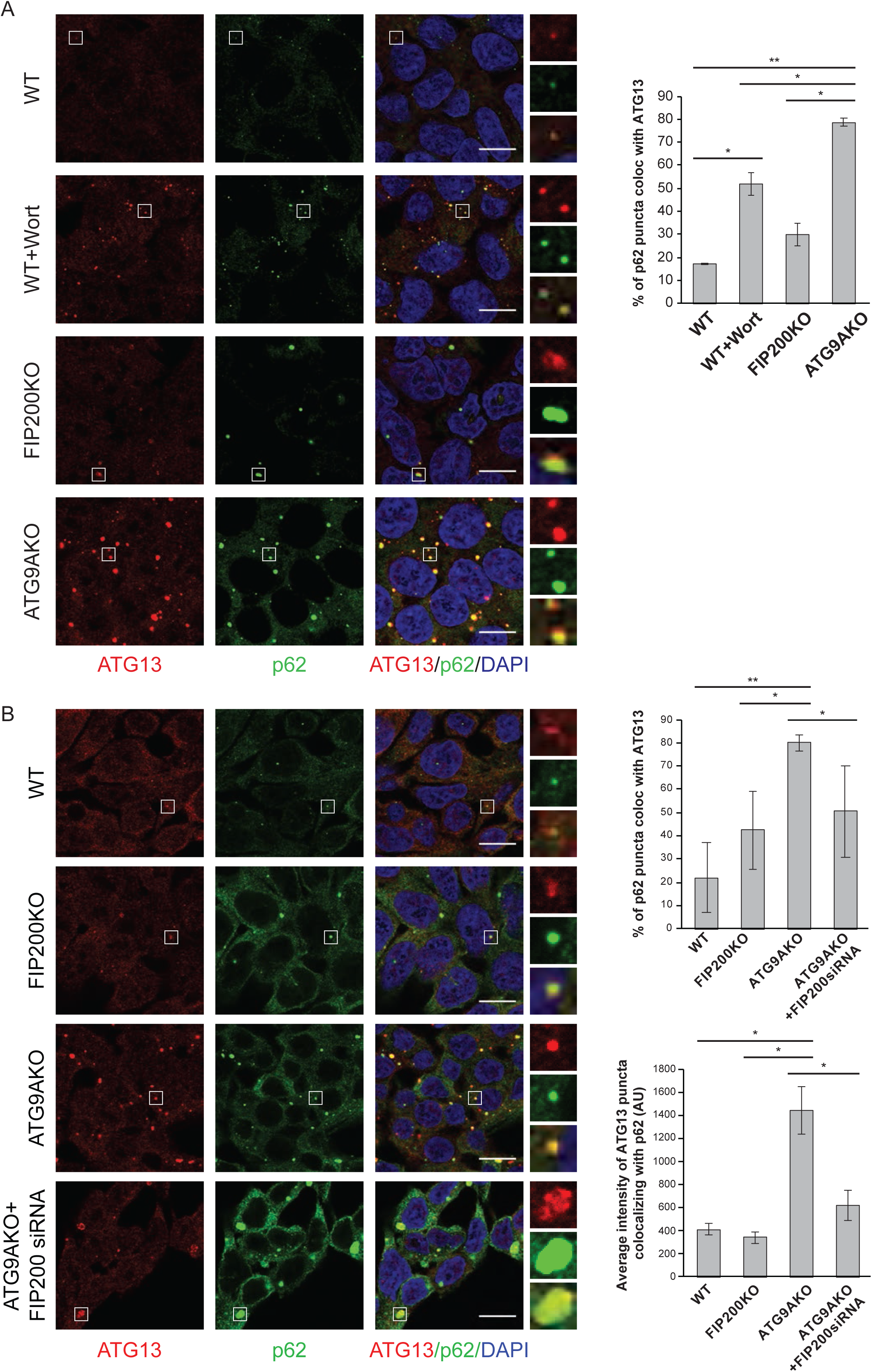
FIP200 is important but not strictly necessary for the recruitment of ATG13 to p62 condensates. (A, B) Colocalization analysis of p62 and ATG13 in HAP1 cells. WT, FIP200 KO and ATG9A KO cells were left untreated or treated with Wortmannin (1µM for 1 h) or with FIP200 siRNA (20 nM for 48 h) as indicated in the figures. Endogenous p62 and ATG13 were detected by immunofluorescence. White boxes represent the magnified areas shown on the right. Scale bar = 10 µm. The percentages of p62 puncta colocalizing with ATG13 are plotted on the right. For panel B, the average signal intensity of ATG13 puncta colocalizing with p62 is also plotted (bottom right). Average values (colocalization percentages or signal intensities) and S.E.M. are shown for n =3. Significances are marked with * when p value ≤ 0,05, ** when p value ≤ 0,01, *** when p value ≤ 0,001.

**Figure 3.**
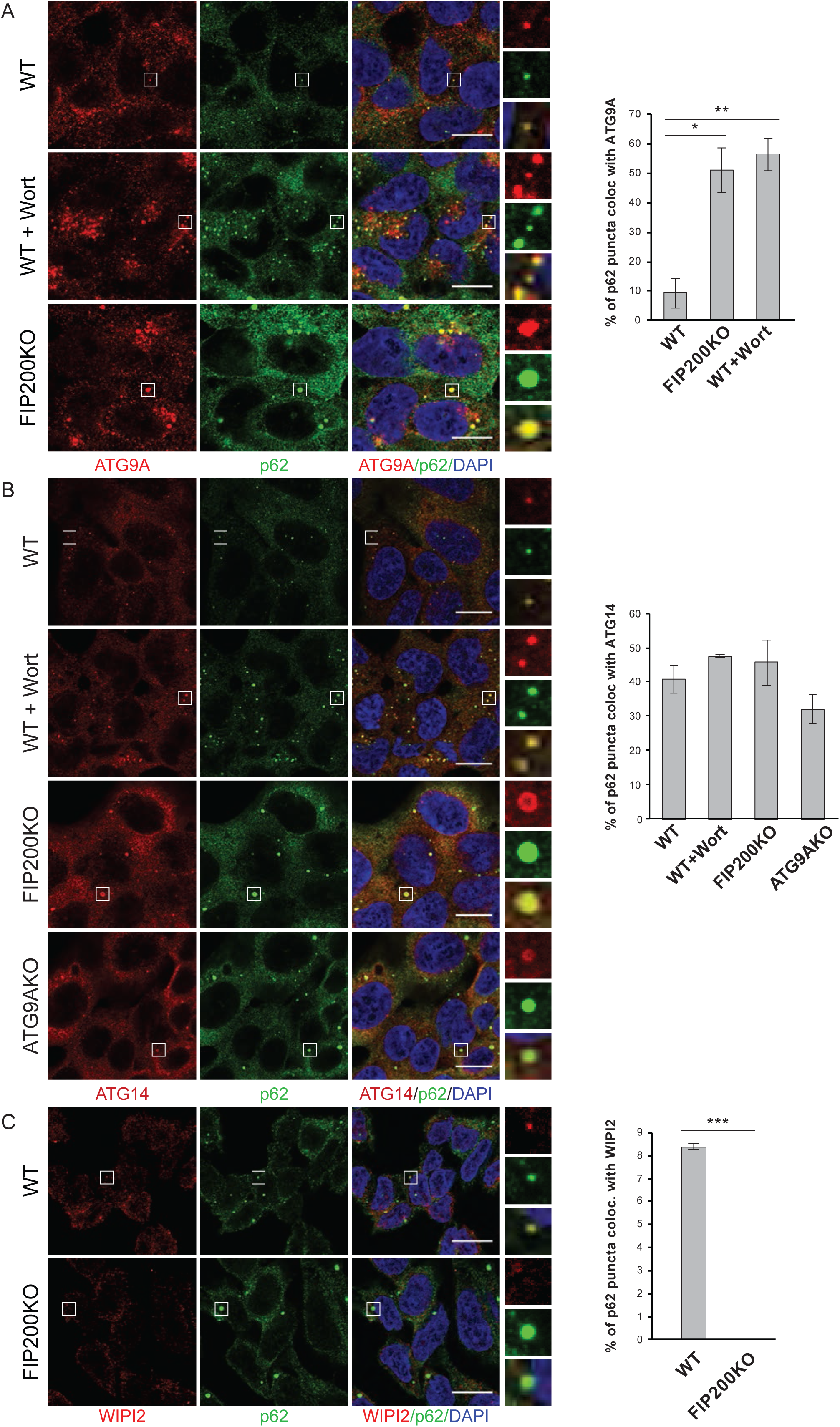
Colocalization of p62 with the components of the autophagy machinery. (A) Colocalization analysis of p62 and ATG9A. HAP1 WT, and FIP200 KO cells were left untreated or treated with Wortmannin (1 µM for 1 h). p62 and ATG9A were detected by immunofluorescence. White boxes represent the magnified areas shown on the right. Scale bar = 10 µm. The percentages of p62 puncta colocalizing with ATG9A are plotted on the right. Average colocalization percentages and S.E.M. for n = 3 are shown. Significances are marked with * when p value ≤ 0,05, ** when p value ≤ 0,01, *** when p value ≤ 0,001. (B) Colocalization analysis of p62 and ATG14. HAP1 WT, FIP200 KO and ATG9A KO cells were left untreated or treated with Wortmannin (1 µM for 1 h). p62 and ATG14 were detected by immunofluorescence. White boxes represent the magnified areas shown on the right. Scale bar = 10 µm. The number (%) of p62 puncta colocalizing with ATG14 is plotted on the right. Average percentages of colocalization and S.E.M. for n = 3 are shown. (C) p62 and WIPI2 colocalization analysis. p62 and WIPI2 were detected by immunofluorescence in HAP1 WT and FIP200 KO cells. White boxes indicate the magnified areas shown on the right. Scale bar = 10 µm. The percentages of p62 puncta colocalizing with WIPI2 are plotted on the right. Average percentages of colocalization and S.E.M. for n = 3 are shown. Significances are marked with * when p value ≤ 0,05, ** when p value ≤ 0,01, *** when p value ≤ 0,001.

In summary, FIP200 is important for the recruitment of the ULK1 complex subunits ATG13 and ULK1, but is not necessary for the recruitment of ATG9A vesicles and the PI3KC3-C1 subunit ATG14 to p62 condensates. Surprisingly, despite the presence of all the upstream components at the p62-ubiquitin condensates, the localization of WIPI2 to these structures is completely abolished in cells lacking FIP200 (Figure 3C). Since WIPI2 recruits the E3 to the site of autophagosome formation (Dooley et al., 2014), this is consistent with our previous observation that FIP200 is essential for the localization of the E3 to p62-ubiquitin condensates (Turco et al., 2019). ATG9A was also essential for the recruitment of WIPI2 to the condensates (Figure EV3). The lack of WIPI2 recruitment strongly suggests that despite the presence of all the upstream components including ULK1, PI3KC3-C1 and ATG9A vesicles in FIP200 deficient cells, these factors are not properly activated and/or spatially arranged to catalyze PI3P production for the recruitment of the downstream machinery.

### FIP200 is necessary for ULK1 activation at p62

We hypothesized that lack of PI3P production in FIP200 KO cells might be due to defective ULK1 activation because the ATG14 and Beclin-1 subunits of the PI3KC3-C1 are substrates of the ULK1 kinase. Upon autophagy inducing conditions, such as starvation or mTOR inhibition by rapamycin, phosphorylation of S29 in ATG14 and S14 in Beclin-1 were shown to stimulate the PI3KC3-C1 activity (Park, Jung et al., 2016, Russell, Tian et al., 2013, Wold, Lim et al., 2016). Also VPS34, the catalytic subunit of the PI3KC3-C1, is phosphorylated at S249 by ULK1, although this post-translational modification seems to have no major effects on autophagy (Egan, Chun et al., 2015).

Here we used ATG14 S29 phosphorylation as a readout for ULK1 activity. In agreement with previous reports (Park et al., 2016, Russell et al., 2013) we found that ATG14 S29 is phosphorylated even in absence of starvation, when only basal autophagy is active. This phosphorylation is dependent on ULK1 and FIP200 (Figure 4A, 4B and EV4), but not on ATG9A (Figure 4B). Structural studies of FIP200 N-terminal domain show that it forms a dimer which interacts with only one molecule each of ATG13 and ULK1 (Shi et al., 2020). This peculiar interaction mode suggests that clustering of more FIP200 dimers is necessary to bring together ULK1 molecules, leading to their clustering and subsequent activation by autophosphorylation (Bach, Larance et al., 2011) and potentially protection from counteracting phosphatases. FIP200 might thus be necessary to allow clustering and autoactivation of ULK1. Interestingly, the R1573D mutation in the FIP200 Claw domain detectably reduced S29 phosphorylation (Figure 4A) mirroring its effect on LC3B lipidation (Figure 1A). This suggests that Claw mediated recruitment of FIP200 is responsible for considerable fraction of cargo-induced selective autophagy that can occur in the presence of mTOR signaling (Turco et al., 2019, Vargas et al., 2019).

**Figure 4.**
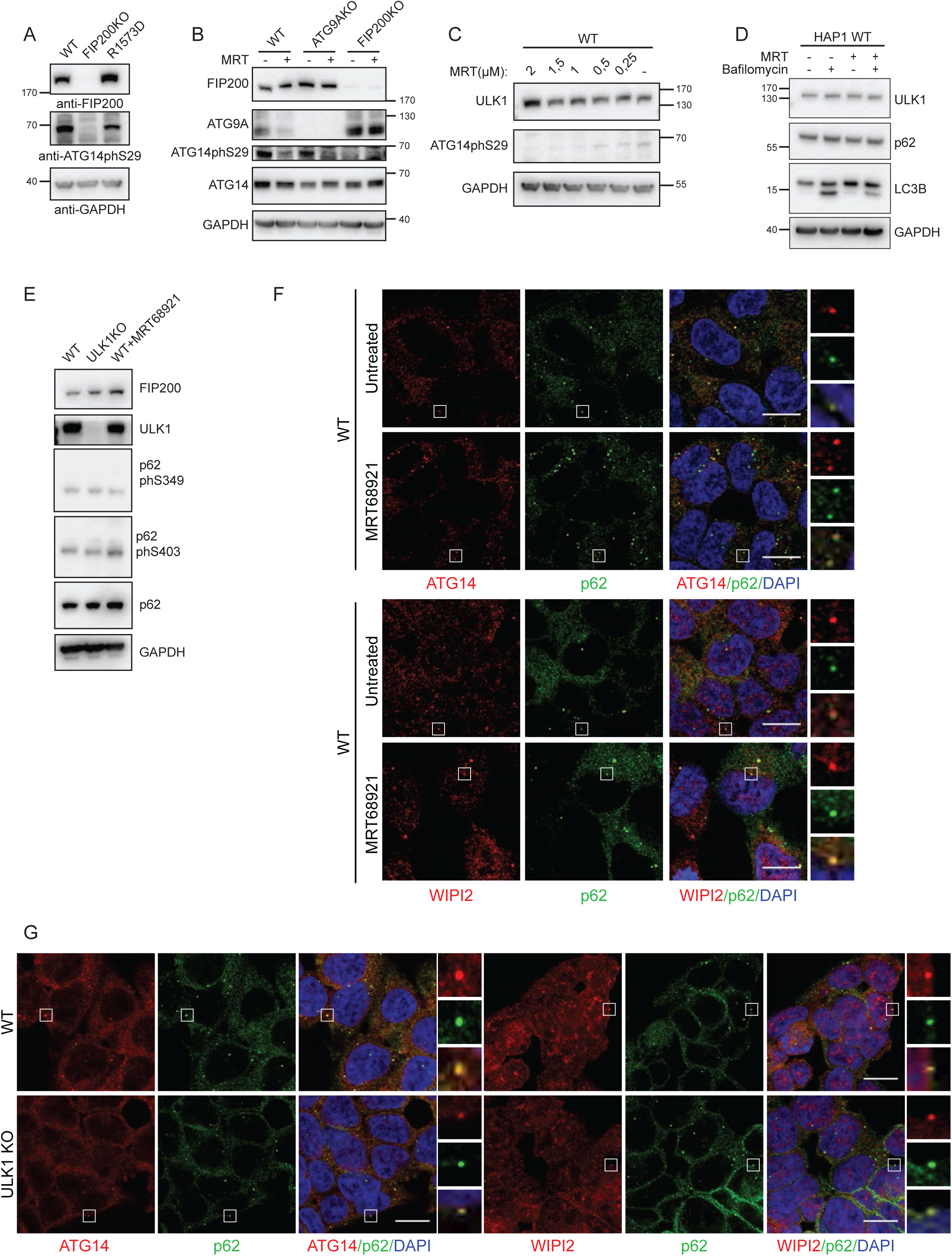
FIP200 is necessary for ULK1 activation. (A) ATG14 phosphorylation in HAP1 cells. Lysates from WT, FIP200 KO and FIP200 R1573D cells were analyzed by western blotting. ATG14 phosphorylation was detected with an ATG14 phS29 antibody. (B) HAP1 WT, ATG9A KO and FIP200 KO cells were left untreated or treated with the ULK1/2 inhibitor MRT68921 (1,5 µM for 2 h). Cell lysates were analyzed by western blotting. FIP200 and ATG9A antibodies were used as a control for the knock out cell lines. Total and phosphorylated ATG14 were detected with anti-ATG14 and anti-ATG14phS29 respectively. (C) Titration of the ULK1/2 inhibitor MRT68921. HAP 1 cells were left untreated or treated with increasing concentration of MRT68921. The ATG14 phS29 antibody was used to monitor ULK1/2 activity. (D) Analysis of autophagy markers upon inhibition of ULK1/2. HAP1 WT cells were left untreated or treated with MRT68921 (1,5 µM for 2 h) and/or Bafilomycin A1 (400 nM for 2 h). Cells lysates were analyzed by western blotting. The ULK1 antibody was used to monitor the protein levels upon inhibition of its kinase activity. p62 levels and LC3B lipidation were detected with the respective antibodies. (E) p62 phosphorylation levels upon inhibition/knock out of ULK1. HAP1 WT and ULK1 KO cells were left untreated or treated with MRT68921 (1,5 µM for 2h) as indicated in the figure. Cell lysates were analyzed by western blotting. FIP200 and ULK1 antibodies were used to monitor the levels of the proteins upon inhibition/deletion of ULK1. Total p62 and phosphorylated p62 were detected with anti-p62 and anti-p62 phS349 or phS403 antibodies. (F, G) Colocalization of p62 with ATG14 and WIPI2 upon inhibition of ULK1/2 (F) or deletion of ULK1 (G). HAP1 WT and ULK1 KO cells were left untreated or treated with MRT68921 (1,5 µM for 2 h). After fixation, p62 and ATG14 or p62 and WIPI2 were detected by immunofluorescence. White boxes represent the magnified areas shown on the right. Scale bar = 10 µm.

If the major role of FIP200 were to activate the ULK1 kinase, which in turn activated the PI3KC3-C1 to produce PI3P, we reasoned that inhibition of the ULK1 activity would phenocopy the effect of FIP200 deletion in the autophagy of p62-ubiquitin condensates. We therefore treated cells with the ULK1/2 inhibitor MRT68921 (Petherick, Conway et al., 2015)(Figure 4C) and followed the degradation of p62 and LC3-II by western blotting. Surprisingly, we did not detect any major defects in the degradation of these proteins, even when we followed the S349 and S403 phosphorylated forms of p62, which are sensitive markers for the selective autophagy of p62-ubiquitin condensates (Figure 4D, E).

Moreover, upon ULK1/2 inhibition both ATG14 and WIPI were efficiently recruited to p62 condensates (Figure 4F), suggesting that the PI3KC3-C1 on the condensates is active and WIPI2 is recruited as a consequence of PI3P formation. To further investigate the role of the ULK1 protein, beside its kinase activity, we used a ULK1 KO cell line. ULK2 was not deleted, since the number of transcript per million (TPM) in HAP 1 WT cells is 0,31, compared to 6,78 TPM for ULK1, according to the data provided by the manufacturer. Similar to ULK1/2 inhibition, ULK1 deletion caused no major phenotype, as shown by colocalization of p62 with ATG14 and WIPI2 (Figure 4G).

We conclude that FIP200 is necessary for ULK1 activation, but, similar to the situation in mitophagy (Zachari, Gudmundsson et al., 2019), ULK1/2 kinase activity and, in this case, the ULK1 protein are dispensable for the autophagic degradation of p62-ubiquitin condensates.

### Super-resolution microscopy of p62 condensates reveals a possible scaffolding function of FIP200

The data presented above suggest that the role of FIP200 may be to organize the upstream components of the autophagy machinery to allow PI3KC3-C1 activity and thus autophagosome formation. Indeed, the elongated structure of FIP200 with the N-terminus engaging in the ULK1 complex formation and the C-terminal end binding to p62 makes it ideal as organizer of the autophagy machinery at the cargo (Shi et al., 2020, Turco et al., 2019). To gain further insights into how FIP200 contributes to the organization of the autophagy machinery we visualized the autophagy machinery at p62 condensates by microscopy. In ATG9A KO cells FIP200 was distributed around p62 condensates, forming a cup-shaped structure. Almost all these structures contained a small FIP200 positive protrusion (Figure 5A). Similar structures were observed for ATG13 (Figure 5B). When we imaged the p62 condensates in ATG9A KO cells by Stimulated Emission Depletion (STED) microscopy, the distribution of FIP200 around the condensates was readily visible, and for each structure one or more FIP200 positive protrusions that were negative for p62 could be identified (Figure 5C, EV5A). Even though they were considerably smaller, the same cup-shaped FIP200 structures around the p62 condensates were also detected in WT cells (Figure 5D). The FIP200 puncta around the condensate probably represent the protrusions seen in ATG9A KO cells. Wortmannin treatment of wild type cells caused accumulation of FIP200 around the condensates, without altering its spatial distribution (Figure 5D). We then imaged ATG13 and p62 in ATG9A KO and FIP200 KO cells. ATG13 formed round structures with lateral protrusion at p62 condensates in ATG9A KO cells (Figure 5E, EV5B). By contrast, in FIP200 KO cells ATG13 clustering in proximity of p62 was still detectable but the spatial organization observed in ATG9A KO cells was completely lost (Figure 5E, EV5B). We made a similar observation for ATG9A, which was scattered at the condensates in the absence of FIP200 (Figure 5F) but formed foci in Wortmannin treated cells. FIP200 and ATG9A also showed a high degree of colocalization in Wortmannin treated cells (Figure 5G). Surprisingly, FIP200 seems to be dispensable for the correct localization of ATG14 at the p62 condensates (Figure 5H).

**Figure 5.**
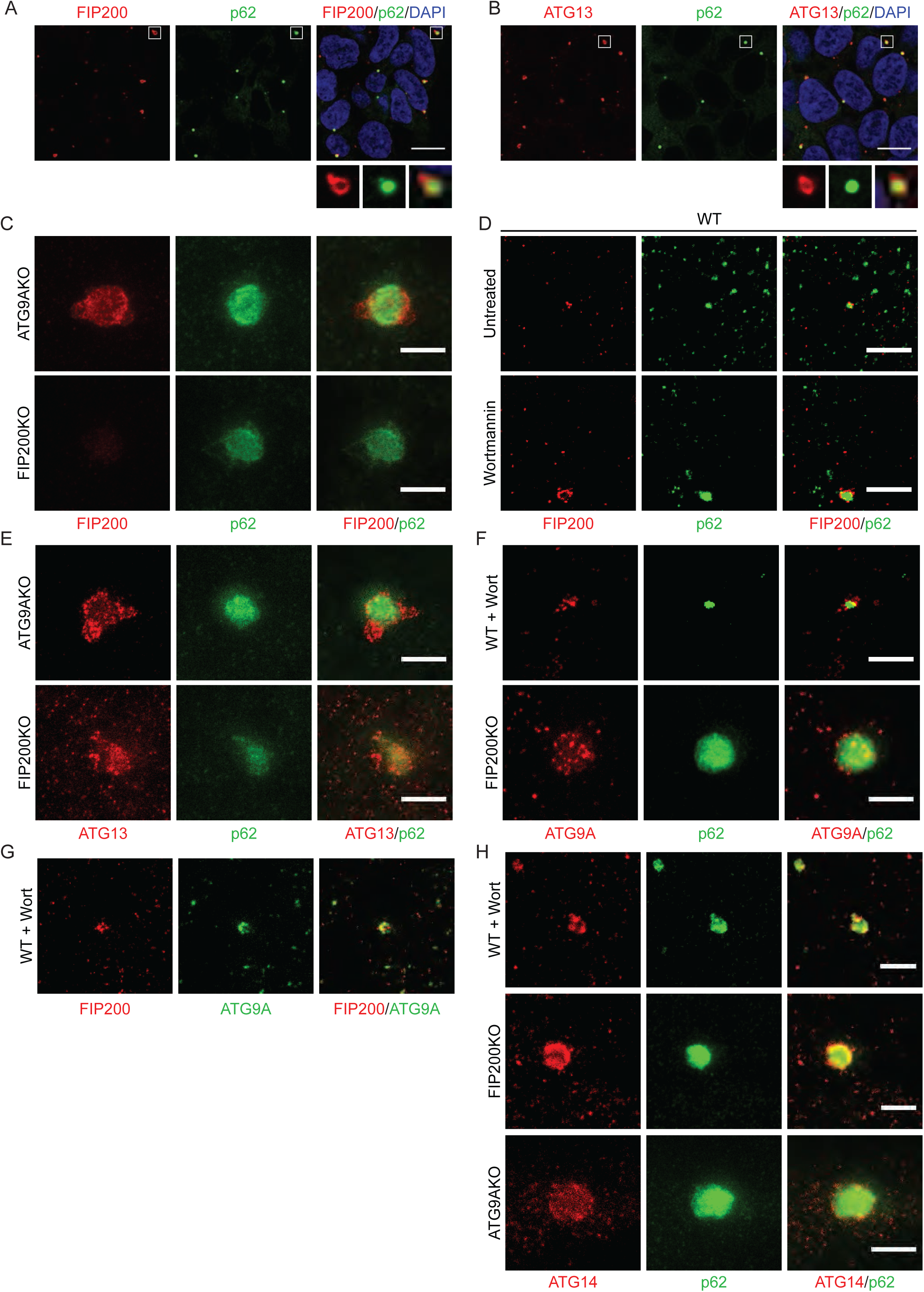
Super-resolution STED microscopy of the autophagy machinery at p62 condensates. (A, B) p62 condensates visualized by confocal microscopy. p62 and FIP200 (A) or p62 and ATG13 (B) were detected by immunofluorescence in HAP1 ATG9A KO cells. White boxes indicate the magnified areas shown under the merged images. Scale bar = 10 µm. (C, D, E, F, G and H) STED microscopy of the autophagy machinery at p62 condensates. Scale bar = 1 µm. In C p62 and FIP200 were detected by immunofluorescence in ATG9A KO and FIP200 KO cells. In D p62 and FIP200 were detected by immunofluorescence in WT cells left untreated or treated with Wortmannin (1 µM for 2 h). In E p62 and ATG13 were detected in ATG9A KO and FIP200 KO cells by immunofluorescence. In F p62 and ATG9A were immuno-stained in WT cells treated with Wortmannin (1 µM for 2h), and FIP200 KO cells. In G immunofluorescence staining of ATG9A and FIP200 was performed in WT cells treated with Wortmannin (1µM for 2 h). In H p62 and ATG14 were detected in WT cells treated with Wortmannin (1µM for 2 h), in FIP200 KO and ATG9A KO cells by immunofluorescence.

In conclusion, although the components of the autophagy machinery can be recruited to p62 condensates, at least to some extent, independently of FIP200, the spatial organization of ATG13 and ULK1 as well as ATG9A was disturbed upon FIP200 deletion.

The arrangement of FIP200 around p62 condensates is reminiscent of the localization of FIP200 and ATG13 in starvation induced autophagy (Karanasios et al., 2016; Kenny et al., 2019) and the frequently observed FIP200 protrusions might represent the sites where the autophagy machinery establishes contacts with the ER (Karanasios et al., 2016, Kenny et al., 2019). FIP200 might thus spatially organize ATG9A vesicles and the ULK1 complex allowing these components to come in contact with the PI3KC3-C1 for subsequent PI3P production. PI3P in turn attracts WIPI2 and the LC3 lipidation machinery for productive autophagosome formation. Unexpectedly, this process seems not to be dependent on the ULK1/2 kinase activity and the ULK1 protein.

In the future, analysis of the ultrastructure of autophagy initiation complexes is likely to provide important insights in the precise relationships between their components.

## Materials and Methods

### Cell culture

HAP 1 cell lines were purchased from Horizon Discovery (https://horizondiscovery.com/en/products/gene-editing/cell-line-models/PIFs/Human-HAP1-Knockout-Cell-Lines) and cultivated in Iscove’s Modified Dulbecco Medium (IMDM), supplemented with 10% Fetal Bovine Serum (Thermo Fisher Scientific), 5,000U/ml Pennicillin/Streptomycin (GIBCO – Thermo Fisher). Cells were cultivated at 37°C in humidified 5% CO_2_ atmosphere.

### Generation of FIP200 R1573D mutant cell line

Mutagenesis of endogenous FIP200 in HAP1 cells by CRISPR/Cas9 was performed using Cas9 nuclease and a dsDNA repair template containing the R1573D mutation. A guide RNA (ACAGATTTAAAGTTCCTTTGGGG) was designed around the R1573 residue of FIP200 using the CRISPOR web tool (https://zlab.bio/guide-design-resources - (Hsu, Scott et al., 2013) and cloned into pSp-GFP-Cas9 plasmid using the BbsI restriction site. For the dsDNA repair template, a 2kb homology region spanning 1kb upstream and downstream of the mutation site was amplified from HAP1 cDNA (primers: ctattGGATCCTCTGGCAGTTATGTTTC; ataagCATATGCACACTTCCCAGCAATC) and cloned into pUC19 plasmid using BamHI/NdeI restriction sites. The R1573D mutation was inserted in the plasmid using Round the Horn PCR mutagenesis. Then, a second cycle of Round the Horn PCR was used to mutate the PAM sequence and to insert a silent BstNI restriction site in proximity of the mutation, to be used for clones’ screening. 40% confluent HAP 1 WT cells (seeded in a 10-cm dish) were co-transfected with the two plasmids described above (5 µg each) using 30 µl Fugene6 transfection reagent (Promega) in OptiMEM medium. 24h after transfection GFP-positive cells were sorted by Fluorescence Activated Cell Sorting (FACS), plated in 10 cm dishes (200000 cells/dish) and let grow until confluent. Single cells were then sorted into 96-well plates for clonal selection. Clones were screened by BstNI restriction digestion of a 500bp PCR product surrounding the mutation site. FIP200 expression levels and the levels of other autophagy-related genes (p62, LC3B) were analyzed by immunoblot and compared to WT HAP1 cells.

### Cell treatment with siRNA and drugs

For siRNA treatment cells were seeded in a 6-well plate (210000 cells/well) and let grow for 24h. Then they were transfected with siRNA (20 nM final concentration) using Lipofectamin RNiMax (Thermo Fisher) in OptiMEM medium. After 48h cells were harvested and processed for further analysis.

Drug treatments were added to the cells before harvesting or fixation according to the following conditions: Bafilomycin A1 – 400 nM, 2h; Wortmannin – 1µM, 2h; MRT68921 – 1,5 µM, 2h.

### Analysis of cell lysates by Immunoblotting

Cells were harvested with trypsin. Cell pellets were washed with PBS and resuspended in 30 µl lysis buffer (50 mM Tris-HCl pH 7,4, 1mM EGTA, 1mM EDTA, 1% Triton X-100, 0,27M sucrose, 1mM DTT, cOmplete EDTA-free protease inhibitor cocktail – Roche, 1mM NaF, 20mM β-glycerophosphate, 1mM Na-vanadate). After 20 min incubation on ice, lysates were cleared by centrifugation at 16000xg for 5 min at 4°C and the total protein concentration in the lysates was measured by Bradford protein assay (Bio-Rad). For western blot analysis 25 µg of each lysate was boiled for 5 min at 98°C (for the detection of LC3B and ATG9A samples were heated at 60°C for 10 min) and resolved by SDS-PAGE. Proteins were then transferred to PVFD membrane by wet blot at 120V for 90 min. Membrane were blocked for 1h at RT with Blocking buffer (3% Non-fat Dry Milk in TBS+0,05% Tween-20). For the detection of p62 ph-S403 and ATG14 ph-S14 membranes were blocked in 5% BSA in TBS+0,05 % Tween-20 (TBST). Primary antibodies were diluted in the respective blocking buffers and incubated with the membrane O/N at 4°C. After 3×15 min washes in TBST, membranes were incubated with Horse Radish Peroxidase conjugated secondary antibody. After 3×15 min washes in TBST signal was detected using Super Signal West Pico/Femto Chemiluminescence Substrate (Thermo Fisher) and a ChemiDoc Touch imaging system (Bio-Rad).

Antibody dilutions: rabbit anti-FIP200 (1:1000, Cell Signaling); mouse anti-p62 (1:3000, BD-Bioscience); mouse anti-LC3B (1:500, Nano-Tools); mouse anti-GAPDH (1:25000, Sigma); rabbit anti-ULK1 H-240 (1:500, Santa Cruz Biotech.); rabbit anti-p62 ph-S349 (1:1000, Cell Signaling); rabbit anti-p62 ph-S403 (1:1000, Cell Signaling); rabbit anti-ATG14 ph-S29 (1:1000, Cell Signaling); rabbit anti-ATG9A (1:1000, Cell Signaling); rabbit anti-ATG14 (1:1000, Cell Signaling); rabbit anti-ATG13 (1:1000, Cell Signaling); rabbit anti-ATG101 (1:1000, Cell Signaling).

Secondary antibodies: goat anti-rabbit (1:10000, Jackson ImmunoResearch); goat anti-mouse (1:10000, Jackson ImmunoResearch).

### Quantification of immunoblotting experiments

Protein bands intensity was measured with ImageJ software (ref). A square was drawn for each gel lane to obtain the lane profile. The area of the peak in the profile was taken as a measure of the band intensity and normalized to the loading control (as specified in the figure legend). Average band intensities and standard error for 3 independent experiments were then plotted.

### Immunocytochemistry

Cells were seeded on ∅12mm high precision glass coverslips (Marienfeld-superior) and let grow for 48 h before fixation in 4% paraformaldehyde (PFA) for 20 min at RT. After fixation coverslips were washes 3 times in PBS and cells were permeabilized with 0,1% Triton X-100 for 5 min at RT. For immunostaining with ULK1 antibody permeabilization was performed in in 0,2% Triton X-100 for 3 min at RT. Samples were then blocked in 1% BSA in PBS for 1 h at RT and subsequently incubated with primary antibody diluted in 1% BSA for 1 h at RT. Coverslips were washed 3 times in PBS and incubated with secondary antibody diluted in 1% BSA for 1h at RT. After three washes in PBS coverslips were mounted on glass slides using DAPI-Fluoromount-G (Southern Biotech). For immunostaining of FIP200 cells were permeabilized with 0,25% Triton X-100 for 15 min at RT. After two washes in PBS coverslips were incubated with primary antibody diluted in 1% BSA for 1 h at 37°C. After 3 x 5 min washes in PBS, samples were incubated with fluorophore conjugated secondary antibody diluted in 1% BSA for 1 h at 37°C and then washed 3 times for 5 min in PBS. Imaging was performed with a Zeiss LSM700 confocal microscope equipped with Plan-Apochromat 63x/1.4 Oil DIC objective. For super-resolution STED microscopy coverslips were mounted with ProLong Glass antifade mountant (Thermo Fisher) and images were taken with an Abberior STEDYCON microscope equipped with alpha Plan-Apochromat 100x/1.46 Oil DIC M27 objective.

Antibody dilutions: mouse anti-p62 (1:100, BD Bioscience – 1:200 for STED microscopy); rabbit anti-p62 (1:500, MBL); rabbit anti-FIP200 (1:200, Cell Signaling); rabbit anti-ATG13 (1:100, Cell Signaling); rabbit anti-ULK1 (1:200, Cell Signaling); goat anti-ATG14 (1:50 Santa Cruz Biotech.); mouse anti-WIPI2 (1:100, Bio-Rad); rabbit anti-ATG9 (1:100, Cell Signaling); rabbit anti-p62 phS349 (1:100, Cell Signaling).

Secondary antibodies: goat anti-mouse AlexaFluor 488 (1:1000, Invitrogen); goat anti-rabbit AlexaFluor 488 (1:1000, Invitrogen); goat anti-mouse AlexaFluor 647 (1:500, Jackson ImmunoResearch); goat anti-rabbit AlexaFluor 647 (1:500, Jackson ImmunoResearch); donkey anti-goat Cy5 (1:200, Jackson ImmunoResearch).

Secondary antibodies for STED microscopy: goat anti-rabbit STAR RED (1:100, Abberior); goat anti-mouse STAR RED (1:200, Abberior); goat anti-mouse AlexaFluor 594 (1:2000, Invitrogen); goat anti-rabbit AlexaFluor 594 (1:2000, Invitrogen); donkey anti-goat AlexaFluor 594 (1:1000, Invitrogen).

### Count of p62 puncta and colocalization analysis

Analysis of the microscopy images was performed using ImageJ. To count and analyze p62 puncta a threshold was applied to the images and the accuracy of the threshold value was validated manually. Then the “Analyze Particles” function was used to count p62 puncta and obtain information on their average size. The number of puncta/cell and their average size was displayed as a mean value from 3 independent experiments and the standard error was calculated. For colocalization analysis images were first thresholded, then puncta in both channels were identified with the “Analyze particle” function and saved in the ROI. Then the identified puncta from both channels were visualized in different colors on the same image and the overlapping puncta were counted. The number of colocalizing puncta/cell was displayed as the average value from 3 independent experiment and standard error was calculated.

### Statistical analysis

For the quantifications of microscopy experiments and western blot statistical significance of the observed difference between two samples was established using a 2-samples unpaired t-test. Significant differences were indicated with * when p-value ≤ 0.05, ** when p value ≤ 0.01, *** when p value ≤ 0.001.

## Acknowledgments

We thank the Max Perutz Labs BioOptics Facility for technical support. This work was supported by the ERC grant No.646653 (S.M.) and by the Austrian Science Fund FWF, No. P30401-B21 (S.M.).

## Authors contributions

E.T. and I.F. conducted the experiments; E.T. and S.M designed the experiments and wrote the paper.

---

**FigureEV1.**
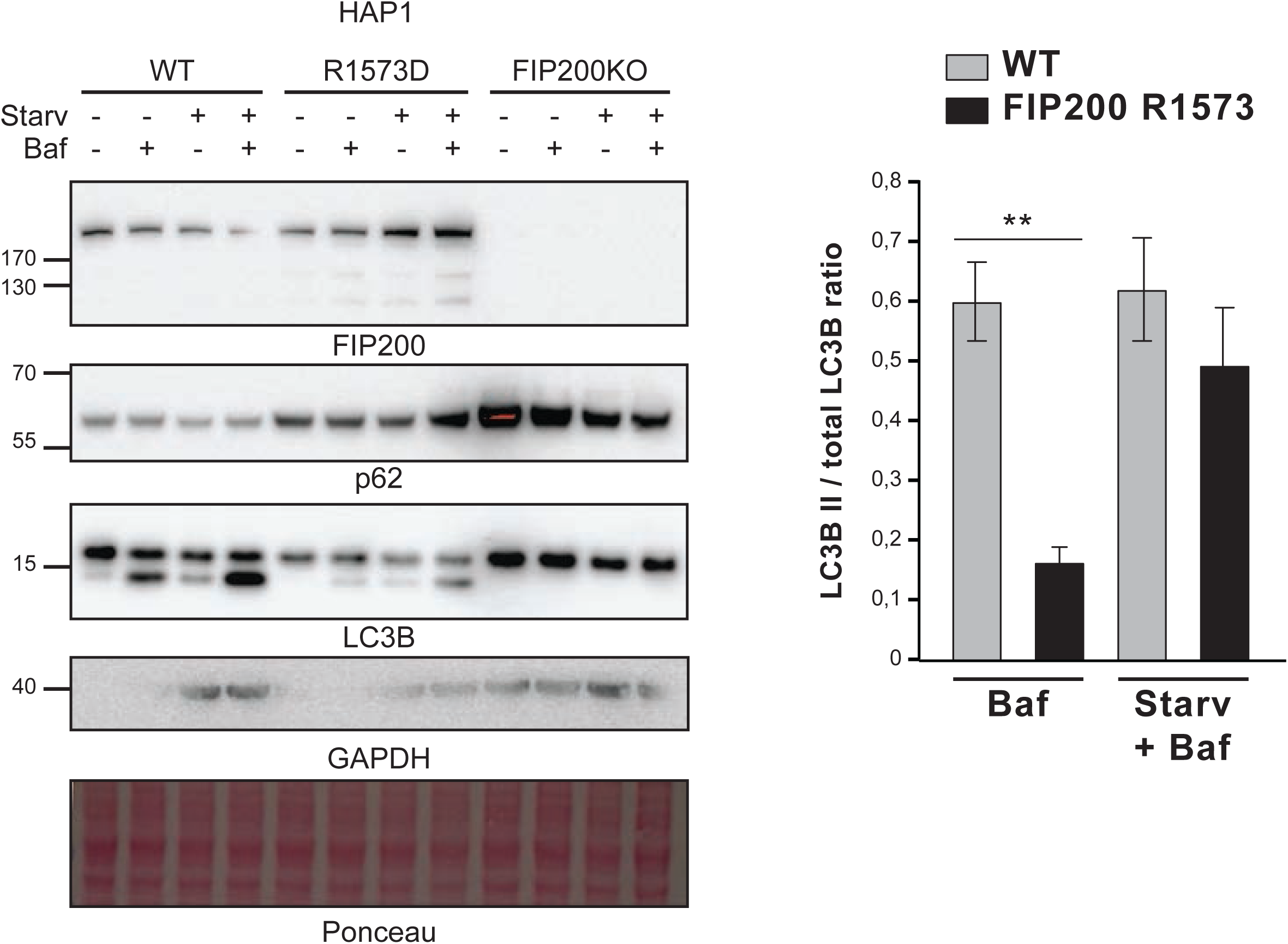
Analysis of p62 and LC3B-II levels in FIP200 KO and FIP200 R1573D mutant cells. WT, FIP200 KO and FIP200 R1573D HAP1 cells were grown in nutrient rich or starvation medium and were left untreated or treated with Bafilomycin A1. Cell lysates were analyzed by western blotting. p62 and LC3B antibodies were used to monitor autophagy. The FIP200 antibody was used to control the levels of FIP200 protein in the mutant cell lines. GAPDH antibody and Ponceau staining of the membrane were used as loading controls. The LC3B-II/total LC3B ratio was calculated and the average ratios from 3 independent experiments are shown in the graph on the right. Error bars represent the S.E.M for n=3. Significances are marked with * when p value ≤ 0,05, ** when p value ≤ 0,01, *** when p value ≤ 0,001.

**FigureEV2.**
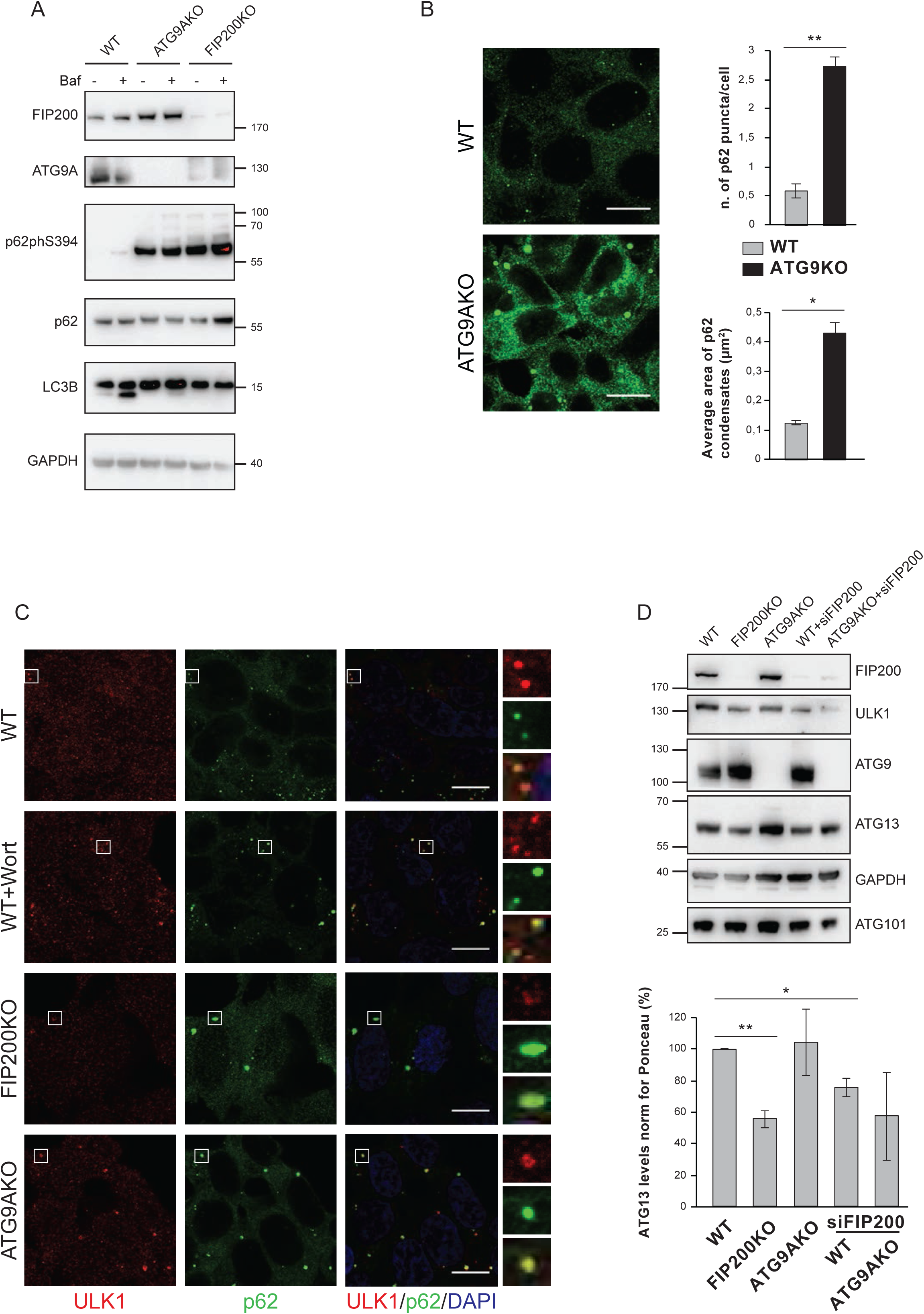
(A) WT, FIP200 KO and ATG9A KO HAP1 cells were left untreated or treated with Bafilomycin A1. Cell lysates were analyzed by western blotting. The FIP200 and ATG9A antibodies were used to control the levels of the proteins in the KO cell lines. Total p62, p62phS349 and LC3B antibodies were used to monitor autophagy (B) p62 condensates were visualized by immunofluorescence staining of p62 in HAP1 WT and ATG9A KO cells. Scale bar = 10 µm. The average number of p62 puncta/cell and their average size in µm^2^ is plotted on the right as the mean value from 3 independent experiments. Error bars represent the S.E.M for n=3. Significances are marked with * when p value ≤ 0,05 and ** when p value ≤ 0,01. (C) p62 colocalization with ULK1 in HAP1 cells. Endogenous p62 and ULK1 were detected by immunofluorescence in WT cells (left untreated or treated with 1 µM Wortmannin for 2h), FIP200 KO and ATG9A KO cells. Scale bars = 10 µm. White boxes represent the magnified areas shown on the right. (D) WT and ATG9A KO HAP1 cells were left untreated or treated with FIP200 siRNA. Cell lysates were then analyzed by western blotting together with FIP200 KO cell lysates. The FIP200 antibody was used to test the efficiency of siRNA treatment. ULK1, ATG13 and ATG101 antibodies were used to monitor the levels of the ULK1 complex components in the KO and siRNA treated cells. The levels of ATG13 protein were measured and plotted as average values from 3 independent experiments. Significances are marked with * when p value ≤ 0,05, ** when p value ≤ 0,01, *** when p value ≤ 0,001.

**FigureEV3.**
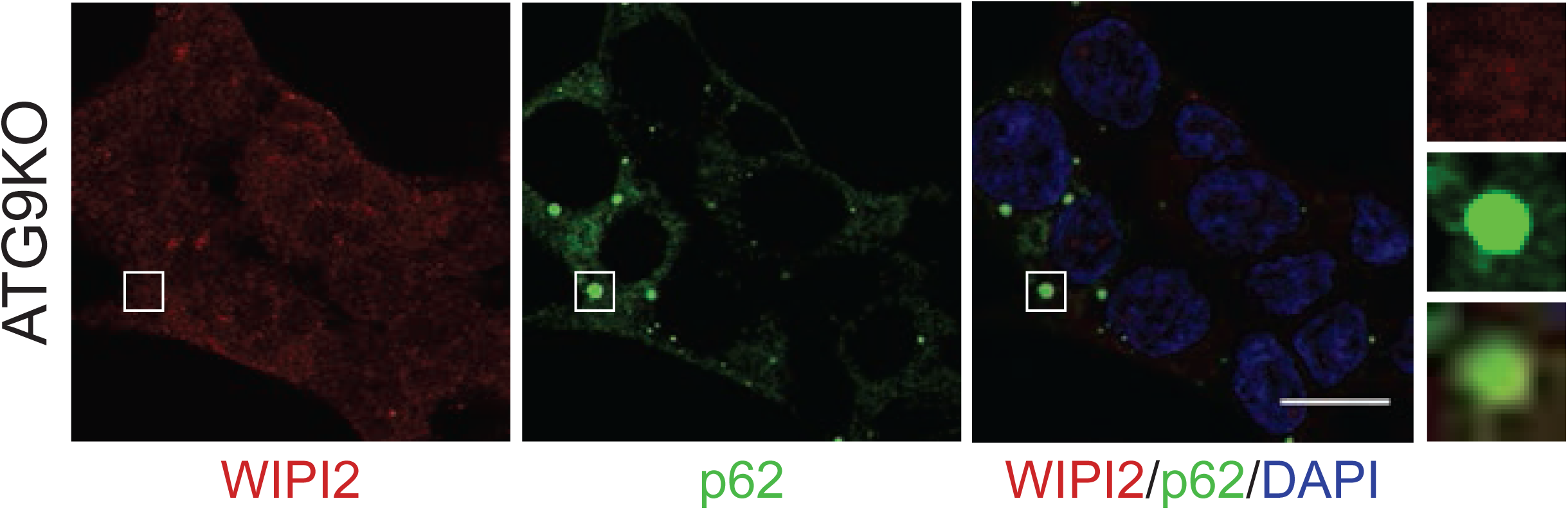
p62 and WIPI2 were detected by immunofluorescence in HAP1 ATG9A KO cells. Scale bar = 10 µm. White boxes indicate the magnified area shown on the right.

**FigureEV4.**
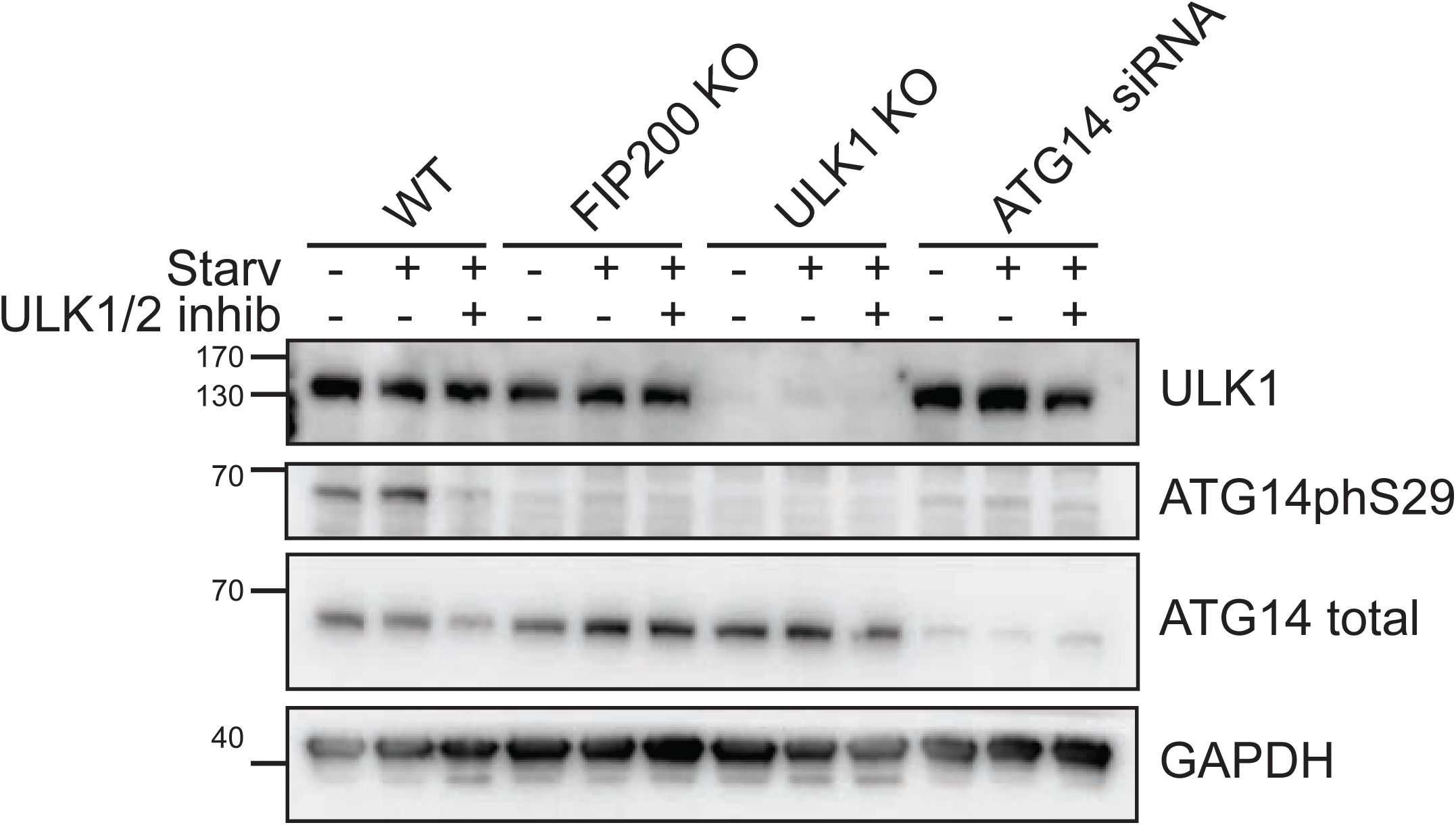
Monitoring ULK1/2 activation in HAP1 cells. WT cells (left untreated or treated with ATG14 siRNA), FIP200 KO and ULK1 KO cells (grown in nutrient rich or starvation medium) were left untreated or treated with ULK1/2 inhibitors. The ATG14 antibody was used to test the efficiency of the siRNA treatment. ULK1 antibody was used to monitor the protein levels upon inhibition of its activity. ATG14phS29 antibody was used to monitor the activity of ULK1/2.

**FigureEV5.**
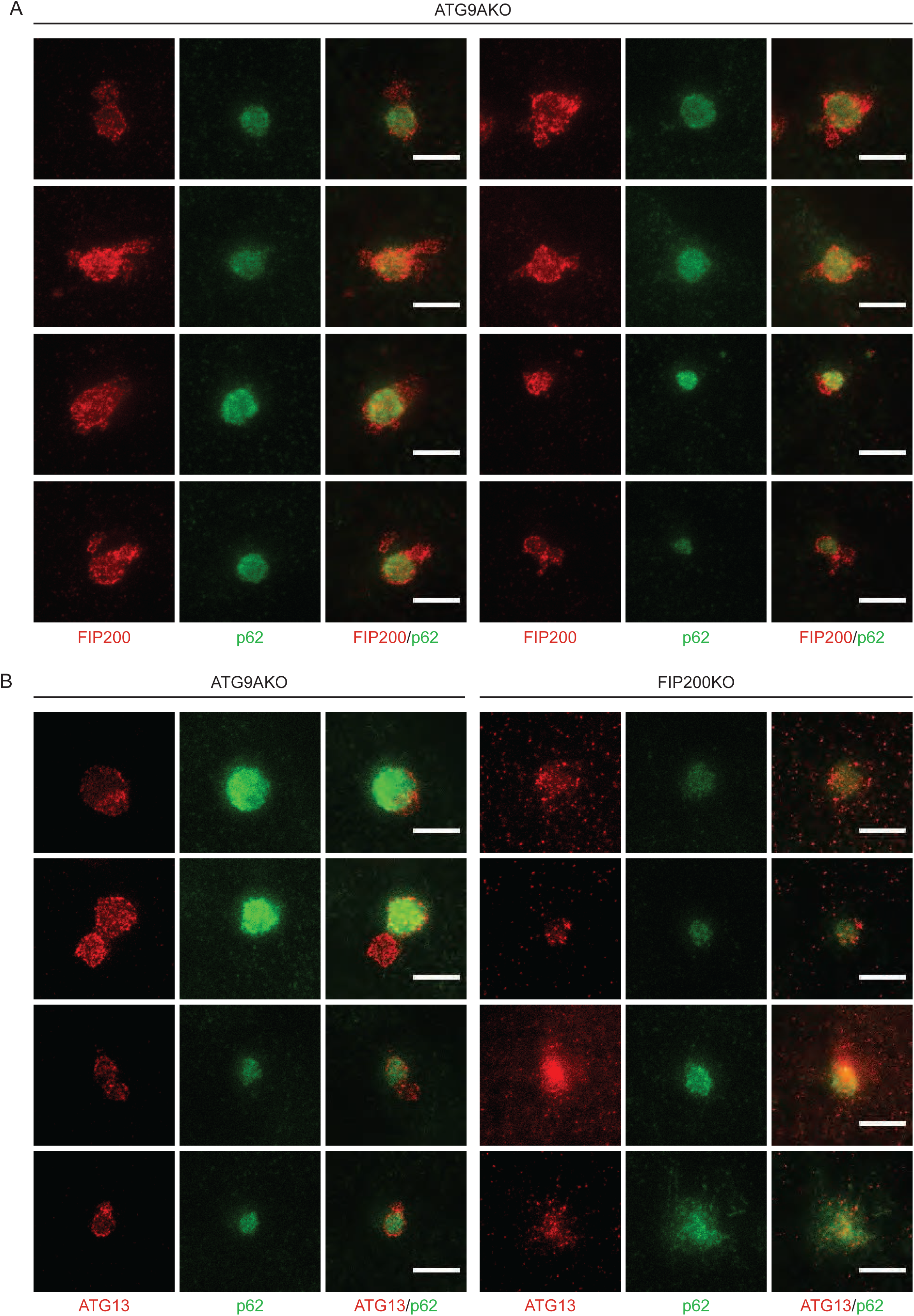
(A, B) STED microscopy of FIP200 (A) and ATG13 (B) at p62 condensates. Endogenous p62 and FIP200 (A) or p62 and ATG13 (B) were detected by immunofluorescence in HAP1 ATG9A KO and FIP200 KO cells. Scale bars = 1 µm.

